# Single cell dynamics of embryonic muscle progenitor cells in zebrafish

**DOI:** 10.1101/396713

**Authors:** Priyanka Sharma, Tyler D. Ruel, Katrinka M. Kocha, Shan Liao, Peng Huang

**Affiliations:** Department of Biochemistry and Molecular Biology, Cumming School of Medicine, Alberta Children’s Hospital Research Institute, University of Calgary, 3330 Hospital Drive, Calgary AB T2N 4N1, Canada; Inflammation Research Network, The Snyder Institute for Chronic Diseases, Department of Microbiology, Immunology and Infectious diseases, Cumming School of Medicine, University of Calgary, 3330 Hospital Drive, Calgary AB T2N 4N1, Canada

**Keywords:** Dermomyotome, Somite, Muscle progenitor cells, In vivo imaging, Extracellular matrix, Zebrafish

## Abstract

Muscle stem cells hold a great therapeutic potential in regenerating damaged muscles. However, the *in vivo* behavior of muscle stem cells during muscle growth and regeneration is still poorly understood. Using zebrafish as a model, we describe the *in vivo* dynamics and function of dermomyotome cells, a population of embryonic muscle progenitor cells. Dermomyotome cells are located in a superficial layer external to muscle fibers and express many extracellular matrix (ECM) genes including *col1a2*. Utilizing a new *col1a2* transgenic line, we show that dermomyotome cells display a ramified morphology with dynamic cellular processes. Cell lineage tracing demonstrates that *col1a2*^*+*^ dermomyotome cells contribute to normal muscle growth as well as muscle injury repair. Combination of live imaging and single cell clonal analysis reveals a highly-choreographed process of muscle regeneration. Activated dermomyotome cells change from the quiescent ramified morphology to a polarized and elongated morphology and generate daughter cells that fuse with existing muscle fibers. Ablation of the dermomyotome severely compromises muscle injury repair. Our work provides a dynamic view of embryonic muscle progenitor cells during zebrafish muscle regeneration.

**Summary statement:** Live imaging and single cell clonal analysis reveal dynamic behaviors of zebrafish embryonic muscle progenitor cells in quiescence and activation.

## INTRODUCTION

Tissue-resident stem cells are crucial for proper organ development and tissue homeostasis. Skeletal muscles possess remarkable ability to regenerate. To harness the power of stem cells for treatment of diseases such as muscular dystrophy, it is critical to understand how muscle progenitor cells behave *in vivo*. In vertebrates, skeletal muscles originate from the somites (Saga and Takeda, 2001). As the embryo develops, the ventral somite forms the sclerotome, generating progenitors of the axial skeleton and tendons, whereas the dorsolateral somite forms the dermomyotome. The dermomyotome further splits to form the dermatome and the myotome, which contribute to the formation of the skin and skeletal muscles, respectively (Christ and Scaal, 2008; Scaal and Christ, 2004). Thus, the dermomyotome contains embryonic muscle progenitor cells required for the initial formation and growth of the musculature. Fate mapping and lineage tracing experiments in mouse and chick have demonstrated that the dermomyotome is also the source of adult muscle stem cells known as satellite cells (Gros et al., 2005; Kassar-Duchossoy et al., 2005; Relaix et al., 2005; Schienda et al., 2006).

Both the dermomyotome (embryonic muscle progenitor cells) and satellite cells (adult muscle stem cells) contribute to muscle fiber formation, and are marked by the expression of the paired box transcription factors *Pax3* and *Pax7* (Dumont et al., 2015; Scaal and Christ, 2004). However, the function and dynamics of satellite cells have been more extensively studied than those of the dermomyotome. Quiescent satellite cells often display a bipolar morphology and reside beneath the basal lamina on the surface of the muscle fiber (Scharner and Zammit, 2011; Webster et al., 2016). The extracellular matrix (ECM) surrounding the satellite cell constitutes its niche and has been implicated in the regulation of satellite cell behavior (Baghdadi et al., 2018; Bentzinger et al., 2013b; Fry et al., 2017; Rayagiri et al., 2018; Tierney et al., 2016; Urciuolo et al., 2013). In the event of muscle injury, “activated” satellite cells undergo proliferation and initiate the myogenic program. The sequential expression of myogenic regulatory factors, including *Myf5*, *MyoD*, and *Myogenin*, results in the differentiation of myoblasts that align and form new syncytial muscle fibers or fuse with existing myofibers. A single transplanted satellite cell is capable of self-renewal and contribute to muscle fibers (Sacco et al., 2008). Conversely, genetic ablation of satellite cells in adult mice completely abolishes injury-induced muscle regeneration (Lepper et al., 2011; Murphy et al., 2011; Sambasivan et al., 2011), demonstrating a critical role of satellite cells in maintaining muscle homeostasis.

It has been challenging to visualize the *in vivo* behavior of satellite cells. Previous work has been mostly inferred from “snapshots” of histological sections or analysis from *in vitro* cultures (Bentzinger et al., 2014; El Fahime et al., 2000; Jockusch and Voigt, 2003; Kuang et al., 2007; Siegel et al., 2009). For example, time-lapse imaging in a 3D myofiber culture system has shown that activated satellite cells are highly dynamic and migrate along the muscle fiber by extending unipolar or bipolar cellular protrusions (Siegel et al., 2009). However, muscle stem cells that are separated from their physiological environment are invariably activated, and *in vitro* approaches therefore do not provide the whole picture of their endogenous behavior. With the advance of intravital imaging, recent work provides the first glimpse of mouse satellite cell behavior *in vivo* (Webster et al., 2016). Quiescent satellite cells are largely immobile, while activated satellite cells proliferate and migrate along the ECM remnants of injured myofibers during regeneration. This work highlights the importance of *in vivo* approaches to study muscle stem cells.

The remarkable regenerative ability and the ease of *in vivo* imaging have made zebrafish a powerful system to study muscle stem cell behavior (Ratnayake and Currie, 2017). The zebrafish dermomyotome, also known as the external cell layer (ECL), is marked by the expression of *pax3* and *pax7*, similar to higher vertebrates (Devoto et al., 2006; Feng et al., 2006; Hammond et al., 2007). During somitogenesis, *pax3/pax7*^*+*^ dermomyotome is generated from the anterior somitic compartment through whole-somite rotation and is thought to generate new muscle fibers during myotome growth (Hollway et al., 2007; Stellabotte et al., 2007). At 4-5 days post-fertilization (dpf), some *pax7*^*+*^ muscle progenitor cells can be observed deep in the myotome between muscle fibers (Seger et al., 2011). Therefore, *pax7* labels at least two populations of muscle progenitor cells in zebrafish embryos: first embryonic muscle progenitor cells in the dermomyotome on the surface of the somite, and later fiber-associated deep myotomal cells, some of which have been shown to be functionally equivalent to mammalian satellite cells (Gurevich et al., 2016). Although *pax7*^*+*^ muscle progenitor cells have been shown to contribute to muscle injury repair (Gurevich et al., 2016; Knappe et al., 2015; Pipalia et al., 2016; Seger et al., 2011), the behavior and contribution of early muscle progenitor cells from the dermomyotome has not been specifically explored due to the lack of specific reporters.

In this study, we developed new transgenic tools and methods to analyze the dynamics of dermomyotome cells at single cell resolution. We identified a number of ECM genes as new markers of dermomyotome cells. Genetic lineage tracing using *col1a2*-based transgenic lines demonstrated that dermomyotome cells contribute not only to embryonic muscle growth but also to injury repair. Using *in vivo* imaging and single cell clonal analysis, we described the dynamics of “quiescent” and “activated” dermomyotome cells. Together, our study provides a dynamic view of embryonic muscle progenitor cells during muscle homeostasis.

## RESULTS

### Extracellular matrix genes are enriched in the dermomyotome

The dermomyotome in zebrafish is traditionally labeled by expression of the muscle progenitor cell marker *pax7* (Devoto et al., 2006; Feng et al., 2006; Hammond et al., 2007). Using double fluorescent in situ hybridization, we identified several extracellular matrix (ECM) genes, including *col1a2* (*collagen 1a2*), *col5a1* (*collagen 5a1*) and *cilp* (*cartilage intermediate layer protein*), that showed co-expression with *pax7* in the dermomyotome on the outer surface of the somite (Fig. 1A). To visualize the dynamics of dermomyotome cells *in vivo*, we generated a *col1a2:Gal4* transgenic line by BAC (bacteria artificial chromosome) recombineering (Fig. 1B). The *col1a2:Gal4* line can be crossed with different UAS lines to label, ablate, or lineage trace dermomyotome cells. Co-labeling using *col1a2* and *kaede* probes in *col1a2:Gal4; UAS:Kaede* (*col1a2*^*Kaede*^ in short) embryos revealed that the *col1a2:Gal4* reporter largely recapitulated the endogenous *col1a2* expression pattern (Fig. S1). Similarly, *col1a2:Gal4; UAS:NTR-mCherry* (*col1a2*^*NTR-mCherry*^ in short) labeled the outer surface of the somite external to *α-actin:GFP*-expressing muscle cells, corresponding to the anatomical location of the dermomyotome (Fig. 1C and Movie 1). The labeling of a few muscle fibers by *col1a2*^*NTR-mCherry*^ suggests that *col1a2*^*+*^ cells contribute to muscle fiber formation. To confirm that the *col1a2:Gal4* line labels the dermomyotome, we performed immunostaining using the anti-Pax7 antibody in *col1a2*^*NTR-mCherry*^ embryos at 2 dpf. Pax7 antibody labels both dermomyotome cells (weaker staining) and xanthophores, neural crest-derived pigment cells (stronger staining). Indeed, all *mCherry*^*+*^ cells on the lateral surface of the somite were weakly *Pax7*^*+*^ (Fig. 1D), indicating that *col1a2*^*NTR-mCherry*^ specifically labels the dermomyotome but not xanthophores. To determine whether the dermomyotome is present in adult zebrafish, we imaged vibratome sections of *col1a2*^*NTR-mCherry*^ fish at 22 mm SL (standard length). We identified three layers of *col1a2*^*+*^ cells external to the muscles, corresponding to scales (the most outer layer), skin (the middle layer), and the presumptive dermomyotome (the most inner layer external to muscles) (Fig. 1E). The presence of a few *mCherry*^*+*^ muscle fibers adjacent to the presumptive dermomyotome suggests that dermomyotome cells contribute to muscle growth in adult zebrafish.

**Figure 1.**
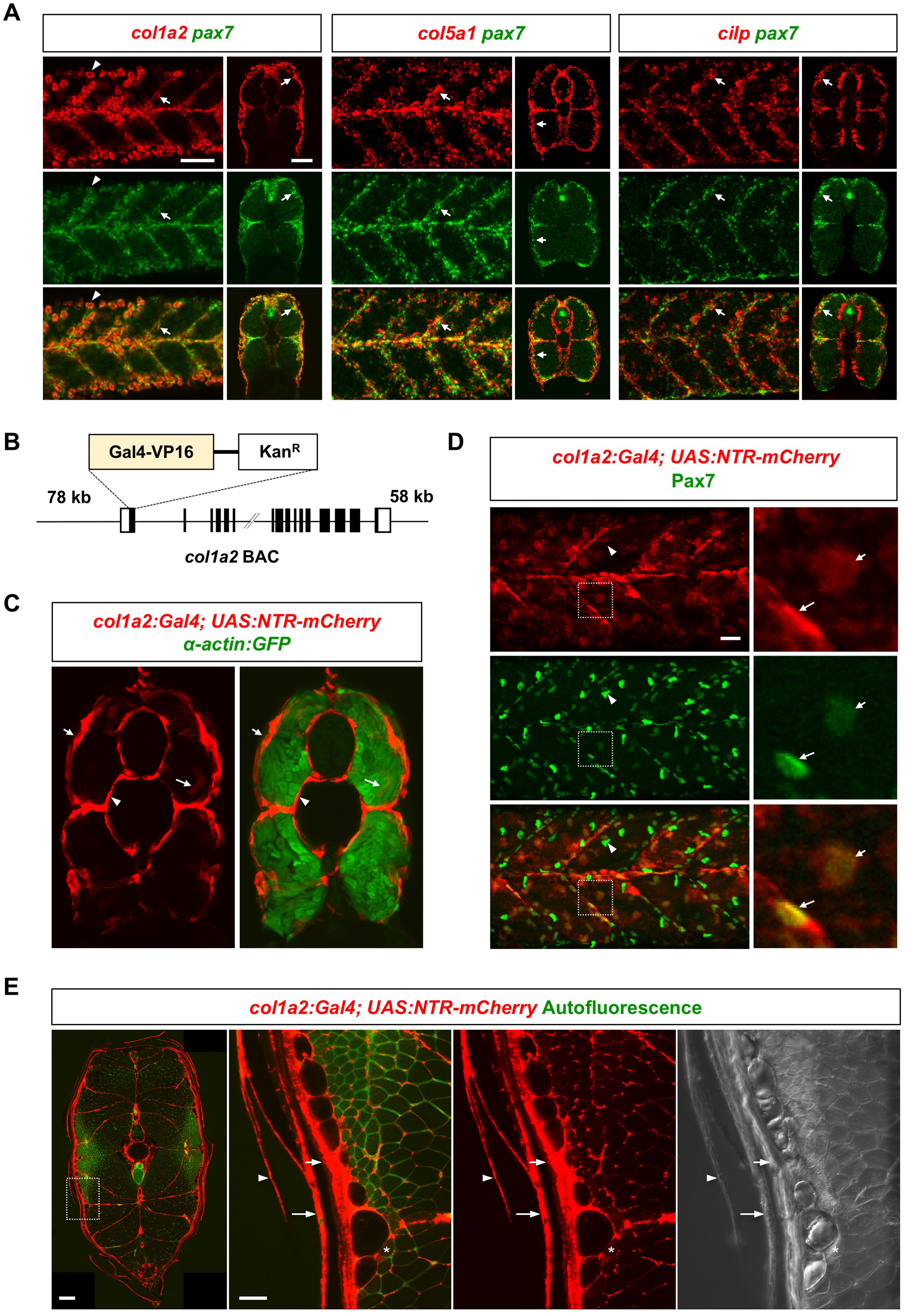
The dermomyotome is marked by the expression of ECM genes. (A) Double fluorescent in situ hybridization showed the co-expression of ECM genes *col1a2*, *col5a1*, and *cilp* (red) with the muscle progenitor cell marker *pax7* (green) in the dermomyotome (arrows) at 3 dpf. Both lateral and transverse views are shown. Note that high-level *col1a2* expression in skin cells bleeds into the red channel (arrowheads). (B) Schematics of the *col1a2:Gal4* BAC reporter. (C) *col1a2*^*NTR-mCherry*^*; α-actin:GFP* fish at 3 dpf showed mCherry expression (red) in the dermomyotome (short arrows) on the surface of muscles, labeled by *α-actin:GFP* (green), cells around the notochord (arrowheads), and occasionally muscle fibers (long arrows). (D) *col1a2*^*NTR-mCherry*^ fish were stained with the anti-Pax7 antibody at 2 dpf. Most *mCherry*^*+*^ dermomyotome cells (red, arrows in expanded views) are co-labeled with Pax7 (weaker staining). Note that Pax7 also labels *mCherry*^*-*^ xanthophores (bean-shaped nuclei with stronger Pax7 staining, arrowheads). (E) Transverse views of *col1a2*^*NTR-mCherry*^ fish at 22 mm SL. Expanded views of the boxed region are shown on the right. mCherry expression (red) can be observed in the presumptive dermomyotome (short arrows), the skin (long arrows), scales (arrowheads), and occasionally muscles (asterisks). The autofluorescence signal (green) is shown to highlight the outline of muscle fibers. Scale bars: 50 µm, except 200 µm in the full view of (E).

### Characterization of the dermomyotome

Using the *col1a2:Gal4* line, we determined the number, distribution, morphology, and dynamics of dermomyotome cells. To quantify the number of dermomyotome cells, we first performed Pax7 antibody staining in *col1a2*^*NTR-mCherry*^ embryos at 2 dpf. On average, there were 25 *Pax7*^*+*^ dermomyotome cells per somite, of which about 87% were also *col1a2*^*+*^ (Fig. 2A), suggesting that the *col1a2*^*NTR-mCherry*^ line labels most dermomyotome cells. The incomplete labeling likely reflects the variegated nature of the Gal4-UAS system (Akitake et al., 2011). Second, *col1a2*^*+*^ dermomyotome cells appeared to distribute evenly to cover the entire surface of the somite (Fig. 1D). About 42% of cells were located along the vertical myoseptum, 43% in between somite boundaries, and 15% along the horizontal myoseptum (Fig. 2B,C). Third, by taking advantage of some highly mosaic *col1a2*^*Kaede*^ embryos, we were able to visualize the morphology of individual dermomyotome cells at 3 dpf (Fig. 2D). They were relatively flat cells sandwiched between the epithelium and muscle fibers. Individual *col1a2*^*+*^ dermomyotome cells appeared to either “float” in between the two somite boundaries or “anchor” their cell bodies along the myoseptum. They always exhibited a remarkable ramified morphology with multipolar lamellipodia-like cellular projections (Fig. 2D). Thin cellular projections (up to 10 µm) extending out of these lamellipodia can often be observed in *col1a2*^*+*^ cells, suggesting potential long range cell-cell communications or the ability to detect distant injuries. By combining two UAS reporters, we visualized the dynamics of dermomyotome cells in *col1a2:Gal4; UAS:NTR-mCherry; UAS:Kaede* embryos (Fig. 2E). The mosaic nature of these transgenes allowed us to label a large number of dermomyotome cells in different colors. Dermomyotome cells appeared to evenly cover the surface of the somite with each cell occupying a non-overlapping territory. Time-lapse imaging showed that after cell division, daughter cells regained the ramified morphology and maintained the similar territory previously occupied by the mother cell (Fig. 2E and Movie 2). Together, our results demonstrate that the new *col1a2:Gal4* driver can be utilized to visualize the dynamics of dermomyotome cells at single cell resolution in zebrafish.

**Figure 2.**
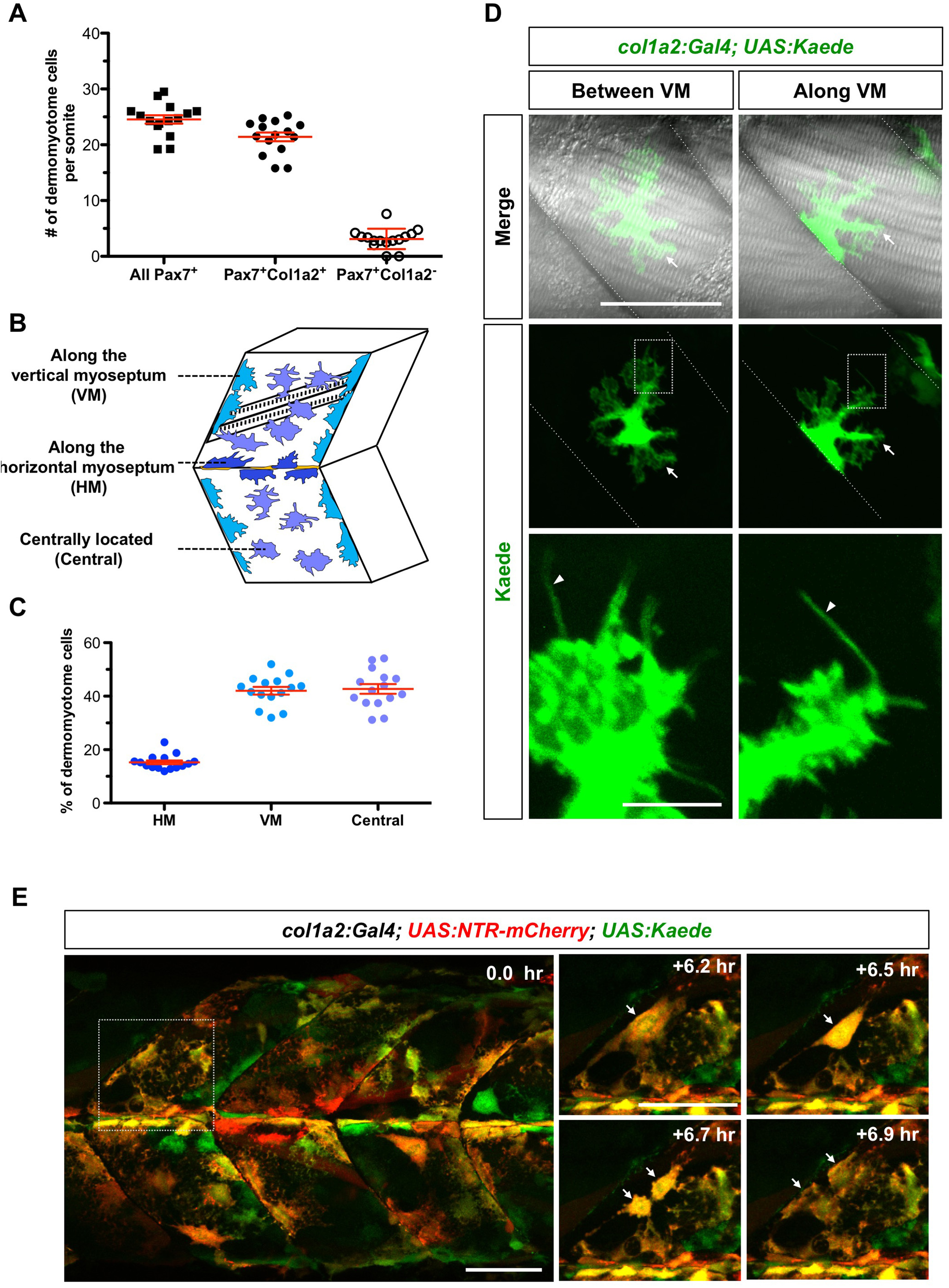
Characterization of *col1a2*^*+*^ dermomyotome. (A) Quantification of dermomyotome cells per somite in *col1a2*^*NTR-mCherry*^ fish stained with anti-Pax7 antibody at 2 dpf. (B) Schematics of dermomyotome cell distribution. (C) Distribution of Pax7^+^Col1a2^+^ dermomyotome cells based on their locations. (D) Mosaic *col1a2*^*Kaede*^ embryos were selected to image single dermomyotome cells. Examples of dermomyotome cells in between vertical myosepta (VM) and along the VM are shown. Dermomyotome cells display ramified morphology with lamelipodia-like structures (arrows) and fine cellular protrusions (arrowheads in expanded views). (E) *col1a2:Gal4; UAS:NTR-mCherry; UAS:Kaede* embryos were imaged at 2 dpf for 7.9 hours. Snapshots from Movie 1 show the division of a dermomyotome cell (arrows). Scale bars: 50 µm, except 20 µm in expanded views in (D).

### Dermomyotome cells generate new muscle fibers during embryonic muscle growth

To determine whether the dermomyotome contributes to muscle growth, we performed two different lineage tracing experiments to follow *col1a2*^*+*^ dermomyotome cells. In the first approach, we took advantage of the photoconvertible fluorescent protein, Kaede (Ando et al., 2002). The normally green-fluorescent Kaede protein (*Kaede*^*green*^) can be photoconverted to a red-fluorescent Kaede protein (*Kaede*^*red*^) by UV light. The perdurance of the *Kaede*^*red*^ protein allows us to trace Kaede-expressing cells for multiple days during development. Briefly, we photoconverted a region of 5-6 somites in *col1a2*^*Kaede*^ embryos at 3 dpf, labeling *col1a2*^*+*^ dermomyotome cells with *Kaede*^*red*^ (Fig. 3A). We then imaged the converted region of the same fish 24 and 48 hours later. New muscle fibers can be easily identified based on their elongated morphology spanning the entire somite between two adjacent vertical myosepta. The emergence of new *Kaede*^*red*^ muscle fibers suggests that *col1a2*^*+*^ dermomyotome cells contribute to new muscle fibers during normal larval development (Fig. 3B).

**Figure 3.**
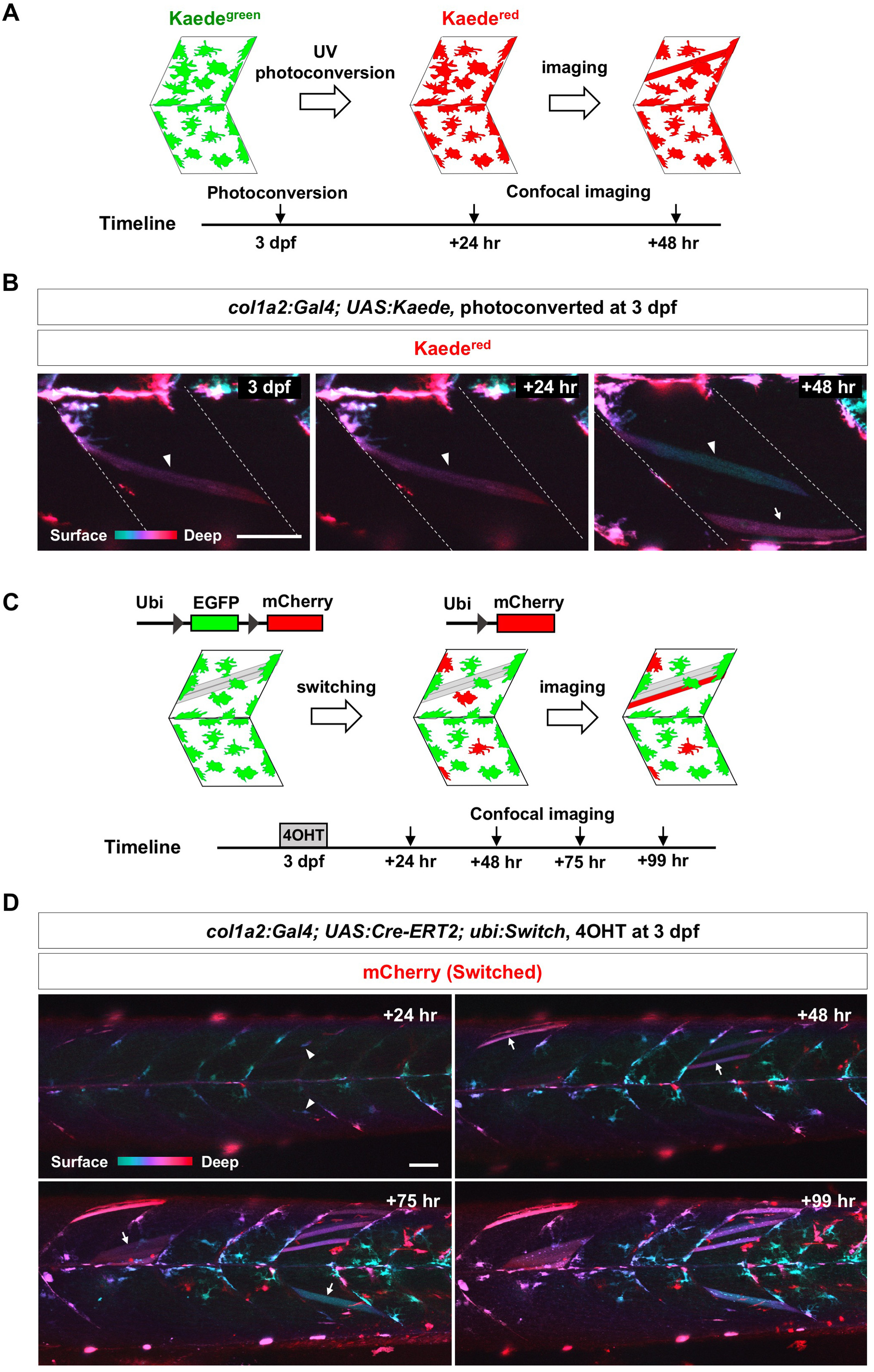
*col1a2*^*+*^ dermomyotome cells contribute to normal muscle growth. (A) Schematics of photoconversion-based lineage tracing. (B) *col1a2*^*Kaede*^ embryos were photoconverted at 3 dpf, and imaged at indicated time points. Color coded depth projections (green corresponds to superficial slices, while red denotes deep slices) of converted *Kaede*^*red*^ showed that new *Kaede*^*red*^ muscle fibers (arrow) emerged at 48-hour post conversion. An existing muscle fiber through all 3 time points is indicated by arrowheads. (C) Schematics of Cre-mediated lineage tracing experiments. (D) *col1a2*^*Cre-ERT^2^*^*; ubi:Switch* embryos were pulsed with 4-OHT for 2 hours at 3 dpf to induce EGFP excision, and imaged for 4 days. Color coded depth projections of the mCherry expression are shown. “Switched” *mCherry*^*+*^ dermomyotome cells (arrowheads) at the “+24 hr” time point generated new *mCherry*^*+*^ muscle fibers (arrows) starting from the “+48 hr” time point. Scale bars: 50 µm.

As a complementary approach, we performed Cre-mediated lineage tracing to determine the fate of *col1a2*^*+*^ dermomyotome cells. We generated the *col1a2:Gal4; UAS:Cre-ERT2* (*col1a2*^*Cre-ERT^2^*^ in short) transgenic line to express tamoxifen-inducible Cre recombinase in dermomyotome cells, and utilized the *ubi:loxP-EGFP-loxP-mCherry* (*ubi:Switch*) line (Mosimann et al., 2011) as the lineage reporter. Induction of Cre activity by 4- hydroxytamoxifen (4-OHT) results in the excision of the EGFP cassette and subsequent expression of the mCherry protein in the cell and its progeny (Fig. 3C). *col1a2*^*Cre-ERT^2^*^*; ubi:Switch* embryos were treated with 4-OHT for 2 hours at 3 dpf and imaged every 24 hours thereafter. At 24 hours after 4-OHT pulse, *col1a2*^*+*^ dermomyotome cells were mosaicly labeled by the mCherry expression, but no muscle fibers were labeled (Fig. 3D). Tracing of *mCherry*^*+*^ dermomyotome cells for 4 consecutive days revealed that new *mCherry*^*+*^ muscle fibers started to emerge at 48 hours post 4-OHT pulse, and increased in number at later time points. Thus, consistent with Kaede-based lineage tracing experiments, this result confirms that dermomyotome cells contribute to the generation of new muscle fibers during muscle homeostasis.

### ECM dynamics during muscle regeneration

We have shown above that *col1a2*^*+*^ dermomyotome cells express muscle progenitor marker *pax7* and contribute to normal muscle growth. To determine whether they also contribute to muscle regeneration, we performed needle injury experiments on muscles of *col1a2*^*NTR-mCherry*^*; α-actin:GFP* embryos (Fig. 4A). Fish were injured within a 1-2 somite area at 3 dpf and imaged every 24 hours for 3 days. At 24 hpi (hours post injury), the injury area, as indicated by the absence of *α-actin:GFP* expression, became substantially smaller than that at 1 hpi. This was accompanied by the emergence of elongated *mCherry*^*+*^ cells in the *GFP*^*-*^ area, suggesting that *col1a2*^*+*^ dermomyotome cells were recruited to the injury site. By 48 hpi, the injury area was completely replaced by newly regenerated muscles, as indicated by higher level of *α-actin:GFP* expression compared to uninjured regions. At 72 hpi, some newly formed muscles were also labeled by *mCherry* expression, suggesting that they were derived from *col1a2*^*+*^ dermomyotome cells. Thus, needle injury experiments demonstrate that small muscle injuries can be quickly repaired within 48 hours in zebrafish and *col1a2*^*+*^ dermomyotome cells are likely the source of new muscle fibers.

**Figure 4.**
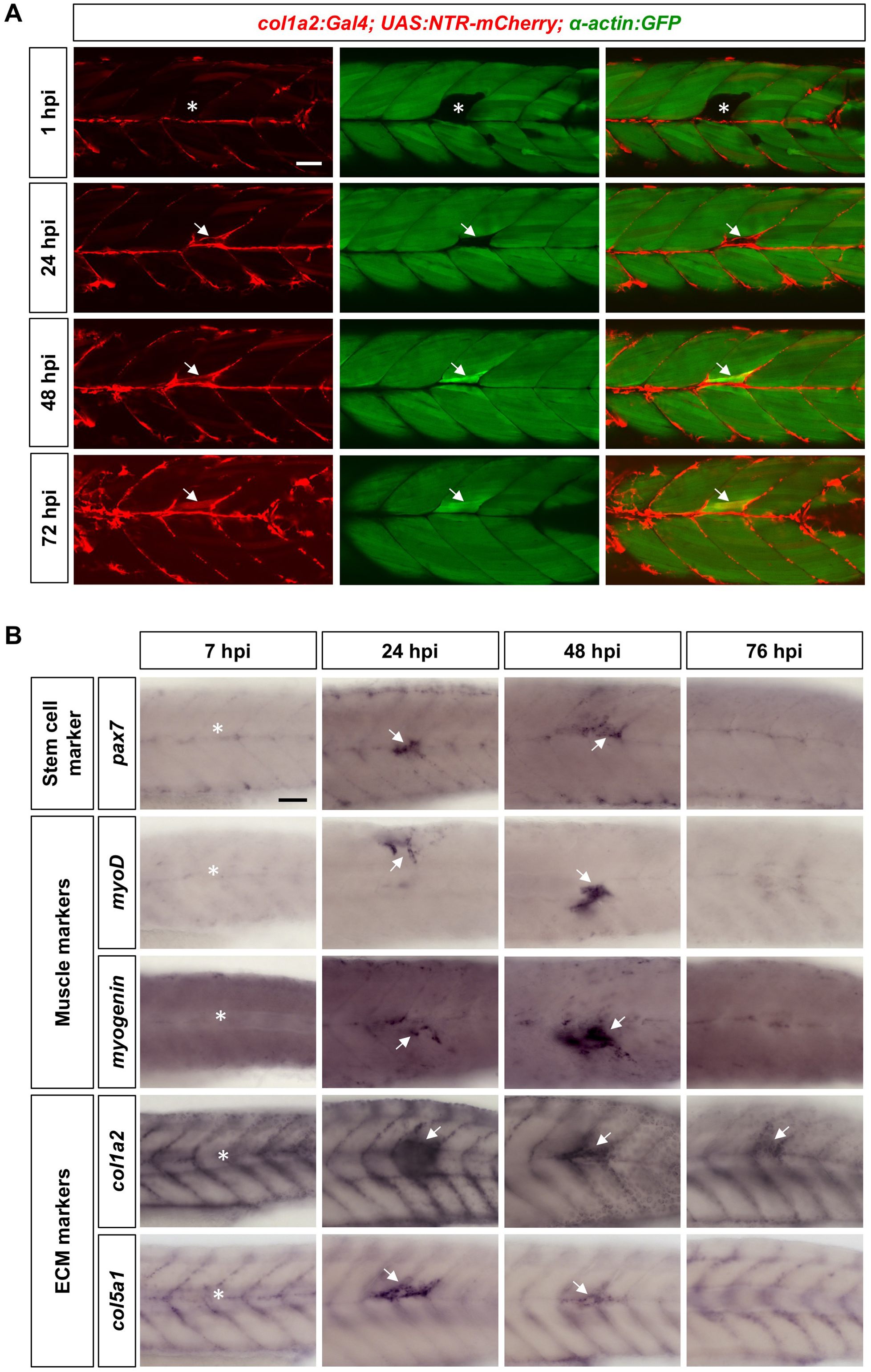
ECM dynamics during muscle injury repair. (A) *col1a2*^*NTR-mCherry*^*; α-actin:GFP* embryos were needle injured at 3 dpf, and imaged at 1, 24, 48, and 72 hpi. Injured muscles (asterisks) can be identified by the lack of *α-actin:GFP* expression (green), while regenerated muscles were marked by elevated *α-actin:GFP* expression. *mCherry*^*+*^ dermomyotome cells (arrows) emerged at the site of injury at 24 hpi, and generated new *mCherry*^*+*^ muscle fibers by 72 hpi. (B) Wild-type embryos were needle stabbed to injure muscles near the end of yolk extension (asterisks) at 3 dpf, fixed at different time points, and stained with the stem cell marker (*pax7*), muscle markers (*myoD* and *myogenin*), and ECM markers (*col1a2* and *col5a1*). All markers showed upregulation at the site of injury starting from 24 hpi (arrows). Scale bars: 50 µm.

We further investigated the expression kinetics of different markers during the entire process of muscle injury repair. Fish were injured by needle stabbing at 3 dpf, and fixed at different time points (7, 24, 48, 76 hpi) for in situ analysis (Fig. 4B). The expression of the muscle stem cell marker *pax7* reached the highest level at the injury site at 24 hpi. The elevated *pax7* expression remained at 48 hpi before returning back to the basal level by 76 hpi. By contrast, myogenic markers (*myoD* and *myogenin*) displayed a kinetics slightly lagging behind *pax7*. Their expression initiated at 24 hpi, reached the peak level at 48 hpi, and returned to the basal level by 76 hpi. This result is consistent with the timing of muscle regeneration from the activation of muscle progenitor cells to the differentiation of new muscle fibers (Fig. 4A). Lastly, analysis of several ECM genes (*col1a2*, *col5a1*, *col1a1a*, *cilp*, *postnb* and *sparc*) revealed similar expression kinetics as *pax7* (Figs 4B and S2). The expression of ECM genes was strongly induced at the injury site at 24 hpi and then gradually declined in the following 48 hours. Together, our results suggest that dermomyotome cells not only are recruited to the injury site to generate new muscle fibers, but also upregulate ECM gene expression perhaps to facilitate the repair.

### Single cell dynamics of dermomyotome cells

We have shown that dermomyotome cells generate new muscle fibers during both normal muscle growth and muscle injury repair. To define the dynamic behavior of individual dermomyotome cells *in vivo*, we performed single cell clonal analysis by taking advantage of the photoconvertible Kaede and the mosaic nature of the *col1a2*^*Kaede*^ transgenic line. Photoconversion of one isolated *Kaede*^*green*^ dermomyotome cell allowed us to visualize its cellular behavior and trace all of its *Kaede*^*red*^ descendants. Briefly, we screened *col1a2*^*Kaede*^ fish at 3 dpf and identified embryos with mosaic labeling of the dermomyotome. Single isolated *Kaede*^*green*^ cells were photoconverted with one cell per somite (3-4 cells per embryo) to ensure accurate cell tracing across multiple time points. We then used the two-photon laser to introduce targeted muscle injury near one *Kaede*^*red*^ cell. *Kaede*^*red*^ dermomyotome cells in laser ablated somites were referred to as cells under injured condition, whereas photoconverted cells in uninjured somites were defined as control cells under wild-type condition. Individual embryos were imaged at 1, 24, 48, and 72 hpi to capture the dynamics of individual converted cells and their descendants (Fig. 5A).

**Figure 5.**
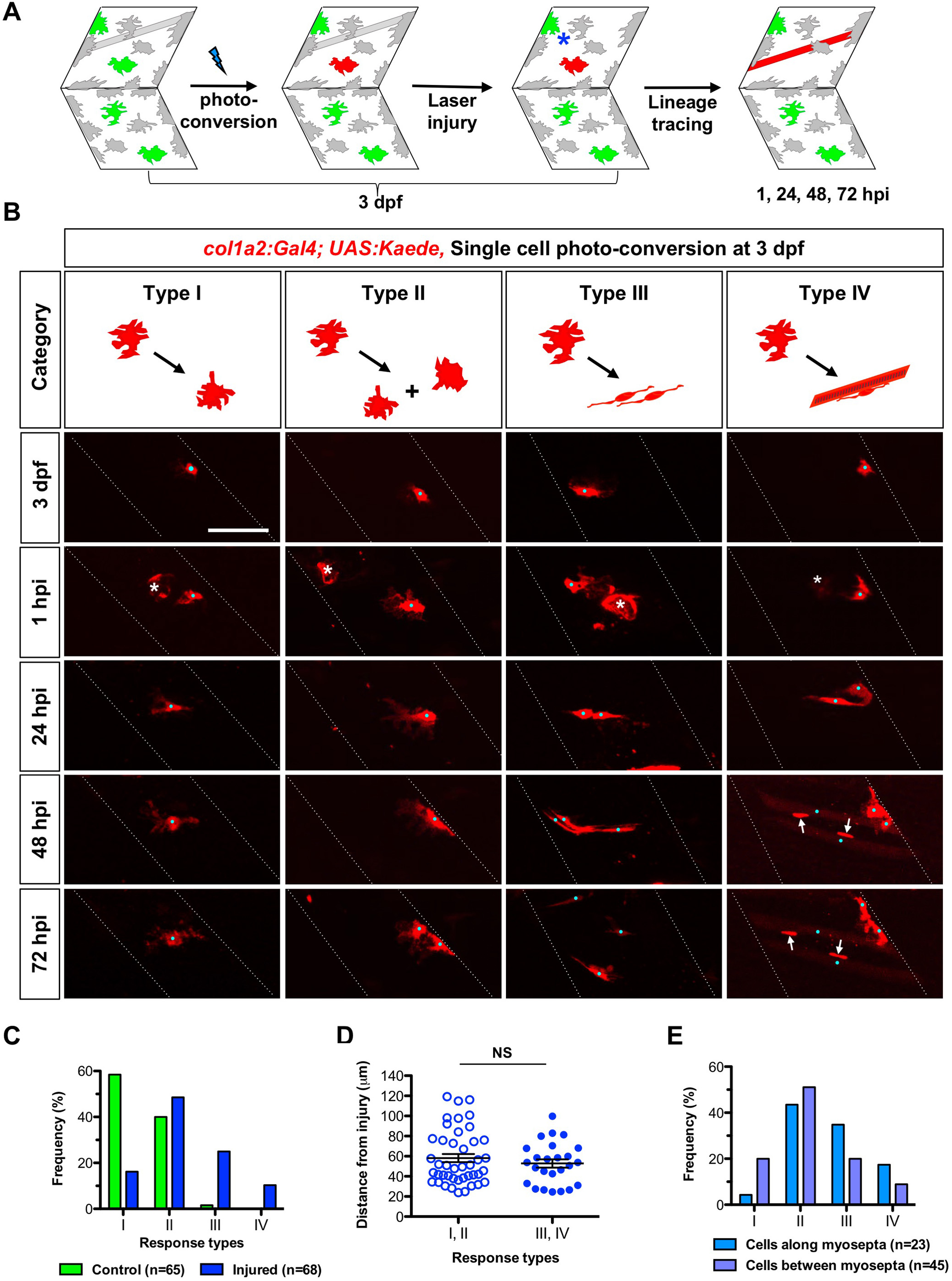
Single cell clonal analysis of dermomyotome cells. (A) Schematics of single cell lineage tracing experiments. Isolated *Kaede*^*green*^ dermomyotome cells in *col1a2*^*Kaede*^ fish at 3 dpf were photoconverted to *Kaede*^*red*^. Immediately after the photoconversion, muscles near the converted *Kaede*^*red*^ cell were damaged by laser ablation (asterisk). The lineage of the *Kaede*^*red*^ cell was inferred by imaging the same region every 24 hours. (B) Single cell lineage tracing in *col1a2*^*Kaede*^ embryos. Four types of responses are represented in cartoons on top with corresponding examples shown at the bottom. Each embryo was imaged at 3 dpf immediately after the photoconversion but before the injury and then at 1, 24, 48, 72 hpi. Note that laser ablation often resulted in elevated autofluorescence at the center of the injury (asterisks). Individual dermomyotome cells and their descendants are marked with cyan dots. The nuclei of newly formed muscle fibers can be identified by high level of *Kaede*^*red*^ expression (arrows). (C) Quantification of response types of dermomyotome cells. (D) Quantification of cell distance from the injury site in different response types. Each point represents one dermomyotome cell under injured condition (68 cells in total). Type I/II (empty dots) and III/IV (solid dots) responses represent quiescent and activated cells, respectively. Data are plotted with mean ± SEM. Statistics: Mann-Whitney *U* test. NS: not significant. (E) Quantification of response types of dermomyotome cells with respect to their initial locations. Cells near myosepta (n=23) are more likely to generate type III/IV responses compared to cells in between myosepta (n=45). Scale bar: 50 µm.

Based on cell behaviors of each clone, we categorized the response of dermomyotome cells into four different categories (Fig. 5B). In type I response, cells did not proliferate and maintained the ramified morphology throughout 72 hours. In type II response, cells underwent one or more cell divisions, but all daughters maintained the ramified morphology, suggesting a quiescent state. By contrast, in type III response, cells generated small elongated cells, which were usually bi-polar with processes extending along muscle fibers. This is markedly distinct from the multipolar ramified morphology of quiescent dermomyotome cells, suggesting an activated state. Lastly, in type IV response, cells first gave rise to small elongated cells similar to those in the type III response, some of which later generated one or more *Kaede*^*red*^ muscle fibers by 72 hpi. As Kaede protein appeared to concentrate in the nuclei of muscle cells (Fig. S3A), new muscle fibers generated from *Kaede*^*red*^ cells can be easily identified based on the stronger *Kaede*^*red*^ signal in the oval-shaped nucleus with a weaker and diffusive signal in the cytoplasm spanning the width of a somite. Antibody staining confirmed that the *Kaede*^*red*^ nucleus of a newly generated muscle no longer expressed Pax7, a feature typical of differentiated muscles, whereas small elongated cells remained *Pax7*^*+*^ (Fig. S3B). Together, under wild-type conditions, dermomyotome cells generated predominantly type I (59%, 38/65 cells) or type II (40%, 26/65 cells) responses, but rarely type III response (2%, 1/65 cells) and never type IV response (Fig. 5C). By contrast, cells under injured conditions exhibited all four types of responses, with a combined 35% (24/68 cells) in the type III and IV categories. Similarly, 24% (16/68 cells) of dermomyotome cells in injured condition generated clones of at least 3 cells compared to only 2% (1/65 cells) in wild-type conditions (Fig. S3C), suggesting an increase in cell proliferation during muscle injury repair. These results suggest that type III/IV behaviors represent the muscle regenerative response of “activated” dermomyotome cells. Since the formation of new muscle fibers was always preceded by small elongated cells (Fig. 5B), the type III response likely represents a transitional phase before the formation of new muscle fibers (type IV response).

Since not all dermomyotome cells would respond to the injury in the same somite (Fig. 5B,C), we asked whether the initial position of the cell influences its behavior. We found that the distance from the injury to the center of the labeled cells does not correlate with the type of response (Fig. 5D). For example, some cells at 24 μm away failed to respond to the injury, while some other cells over 80 μm away became activated. This result suggests that dermomyotome cells can detect and respond to an injury at a long distance away from the cell body. Interestingly, dermomyotome cells located along the myoseptum are more likely to generate a type III or IV response (52%, 12/23 cells) than centrally located cells (29%, 13/45 cells) (Fig. 5E), suggesting that the local niche might influence the behavior of muscle progenitor cells.

### New muscle fibers are predominantly generated by fusion

Our single cell lineage tracing experiments indicate that “activated” dermomyotome cells go through a series of stereotypic phases to generate new muscle fibers (Fig. 5). To further confirm this, we carried out confocal time-lapse imaging to visualize the entire process of muscle regeneration. First, we imaged the “early phase” of muscle repair between 0 to 24 hpi (Fig. 6A). Moderately mosaic *col1a2*^*Kaede*^ embryos were injured by needle stabbing at 59 hpf. Isolated *Kaede*^*green*^ cells were photoconverted to facilitate cell tracking (Fig. 6B and Movie 3). Within the first 24 hours, the “activated” dermomyotome cell started to project polarized cellular processes along muscle fibers. This bi-polar and elongated morphology was maintained even after the cell division. Consistent with our previous observations, new muscle fibers were rarely generated during this time interval. Thus, the “early phase” of muscle regeneration is characterized by the morphological changes and proliferations of “activated” dermomyotome cells. Next, we performed time-lapse imaging of the “late phase” of muscle injury repair. Mosaic *col1a2*^*Kaede*^ embryos were injured by needle stabbing at 3 dpf and imaged from 29 to 48 hpi (Fig. 6C). From a total of 9 movies collected, we observed the generation of 13 new muscle fibers. Remarkably, all 13 fibers were formed in a similar fashion: a small elongated *Kaede*^*+*^ cell at one time point disappeared by the next time point (8-minute intervals), with simultaneous emergence of *Kaede*^*+*^ muscle fiber characterized by *Kaede*^*strong*^ nucleus and *Kaede*^*weak*^ cytoplasm (Fig. 6D). The rapidity of this event suggests that new muscle fibers are formed through cell fusions between a *Kaede*^*+*^ dermomyotome derived cell and an existing non-labeled muscle fiber.

**Figure 6.**
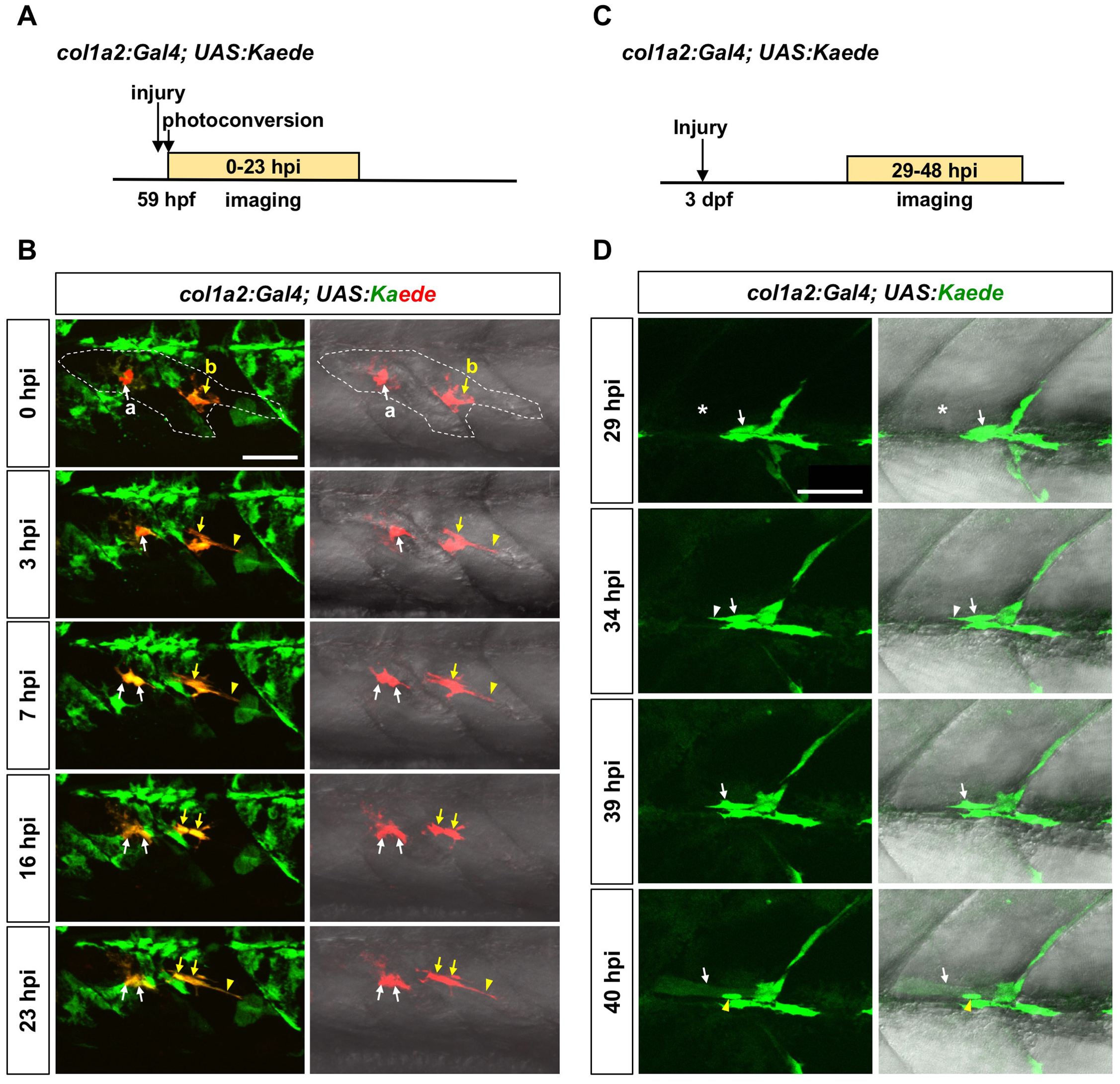
*In vivo* dynamics of dermomyotome cells. (A) Schematic representation of the experiment in (B). *col1a2*^*Kaede*^ embryos were needle injured, photoconverted at 59 hpf, and then imaged from 0 to 23 hpi. (B) Representative snapshots from Movie 3 show the dynamics of 2 photoconverted *Kaede*^*red*^ cells during first 23 hours of regeneration. Both cells were within the muscle injury area, which spanned two somites (outlined by dash lines). Cell “a” (white arrows) maintained the ramified morphology, and divided once at 7 hpi generating two daughter cells with similar morphologies. By contrast, cell “b” (yellow arrows) extended to form an elongated morphology (arrowheads), and divided once at 16 hpi generating two polarized daughter cells. (C) Schematic representation of the experiment in (D). Mosaic *col1a2*^*Kaede*^ embryos were injured at 3 dpf, and imaged from 29 to 48 hpi. (D) A *Kaede*^*+*^ dermomyotome cell (arrows) near the injury site (asterisks) elongated at 34 hpi (white arrowheads), formed protrusions at 39 hpi, and fused with a neighboring muscle fiber at 40 hpi. The newly formed muscle fiber can be visualized by the weak Kaede expression throughout the muscle fiber and the strong Kaede expression in the nucleus (yellow arrowheads). Scale bars: 50 µm.

Since we never observed any *de novo* fiber formation in our time-lapse movies, we asked whether muscles are formed only through cell fusions. To answer this question, we performed genetic lineage tracing in the *col1a2*^*Cre-ERT^2^*^*; ubi:Switch* fish (Fig. 7A). Embryos at 3 dpf were treated with 4-OHT for 3.5 hours to mosaicly label dermomyotome cells. Fish were then injured by needle stabbing and imaged at 75 hpi to quantify newly formed *mCherry*^*+*^ muscle fibers. If a new muscle fiber is generated *de novo*, it would be *mCherry*^*+*^ but *EGFP*^*-*^ due to the excision of the EGFP cassette by Cre-mediated switching. Conversely, if an *mCherry*^*+*^ dermomyotome cell fuses with an already existing muscle fiber (*EGFP*^*+*^), the resulting new muscle fiber will express both mCherry and EGFP (Fig. 7B). In uninjured control embryos, 97% of new muscle fibers (37/38) were formed via cell fusion, whereas only 3% of fibers (1/38) were generated *de novo* (Fig. 7C,D). Interestingly, in injured embryos, *de novo* fiber formation increased slightly to 14% (14/98) at the muscle injury site. Together, this result is consistent with our time-lapse imaging that new muscle fibers are generated predominantly through cell fusion of dermomyotome descendants with existing muscle fibers.

**Figure 7.**
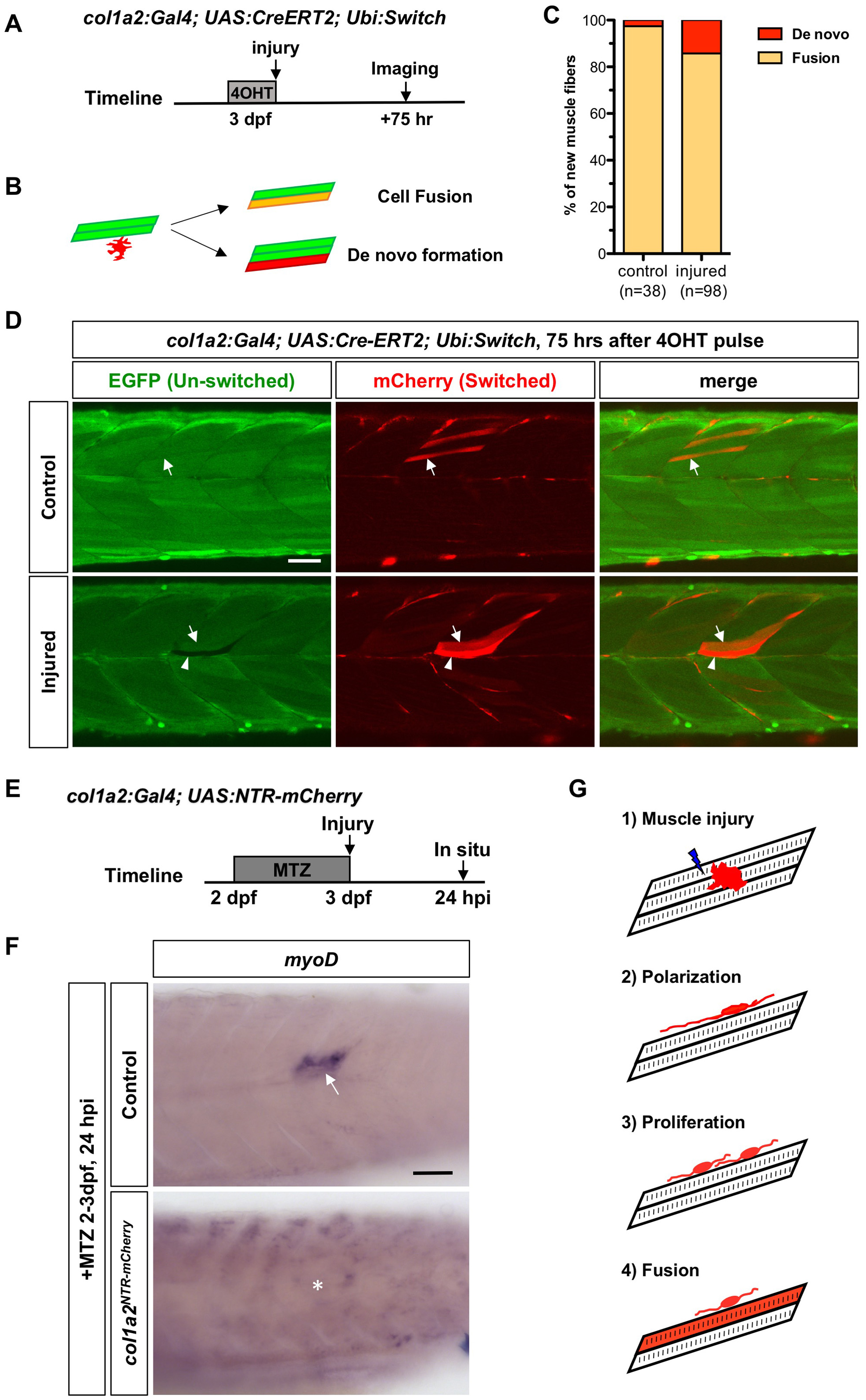
*col1a2*^*+*^ dermomyotome cells generate new muscle fibers primarily by cell fusion. (A) Schematics of lineage tracing experiments. *co1a2*^*Cre-ERT^2^*^*; ubi:Switch* embryos treated with 4-OHT for 3.5 hours at 3 dpf were either needle injured or left uninjured (controls) and imaged 75 hours after the 4-OHT treatment. (B) Two possible modes of new muscle fiber formation. Cell fusion would generate an *mCherry*^*+*^*EGFP*^*+*^ fiber (yellow), whereas *de novo* fiber formation would result in an *mCherry*^*+*^*EGFP*^*-*^ fiber (red). (C) Quantification of two modes of new fiber formation in control and injured embryos. (D) In control embryos, new muscle fibers were formed primarily through cell fusion (arrows), whereas in injured embryos, new fibers were generated by both fusion (arrows) and occasionally *de novo* fiber formation (arrowheads). (E) Schematics of dermomyotome ablation experiment. (F) *col1a2*^*NTR*-mCherry^ or control embryos treated with MTZ at 2-3 dpf were injured at 3 dpf and stained for *myoD* expression at 24 hpi. All control embryos (38/38) displayed significant *myoD* upregulation (arrow), while most ablated embryos (68/88) showed no specific *myoD* induction at the injury site (asterisk). (G) Model of muscle injury repair by dermomyotome cells. Upon muscle injury (1), activated dermomyotome cells transform from the ramified morphology to a highly polarized morphology extending long cellular projections along muscle fibers (2). The activated cell proliferates to generate more elongated daughter cells (3), some of which fuse with an existing muscle fiber to regenerate the damaged region (4). Scale bars: 50 µm.

### Dermomyotome cells are required for effective muscle regeneration

To test whether dermomyotome cells are essential for embryonic muscle injury repair, we ablated *col1a2*^*+*^ dermomyotome cells using the nitroreductase (NTR) based system. The NTR enzyme converts the harmless prodrug metronidazole (MTZ) into a cytotoxic compound that induces rapid cell death of NTR-expressing cells (Curado et al., 2008; Pisharath et al., 2007). *col1a2*^*NTR-mCherry*^ or sibling control embryos were treated with MTZ from 2 to 3 dpf, injured at 3 dpf, and fixed at 24 hpi for in situ analysis (Fig. 7E). In control embryos, *myoD* expression was significantly upregulated at the site of injury (Fig. 7F), suggesting a robust regenerative response. By contrast, most MTZ-treated *col1a2*^*NTR-mCherry*^ embryos showed no specific induction of *myoD* at the injury site. This result suggests that *col1a2*^*+*^ dermomyotome cells are required for an efficient muscle injury repair in early zebrafish embryos.

## DISCUSSION

Using zebrafish as a model, our study provides a dynamic view of embryonic muscle progenitor cells in the dermomyotome. First, dermomyotome cells display a unique ramified morphology expressing many ECM genes. Second, lineage tracing experiments show that dermomyotome cells contribute to normal muscle growth as well as muscle injury repair. Third, dermomyotome cells undergo a series of stereotypical steps to generate new muscle fibers, predominantly by cell fusion with existing fibers.

### Dermomyotome cells are embryonic muscle progenitor cells

Dermomyotome is an evolutionarily conserved structure on the external surface of the myotome present in most vertebrates (Devoto et al., 2006). In zebrafish, the dermomyotome displays remarkable similarities to that of higher vertebrates. Our work and that of others (Devoto et al., 2006; Feng et al., 2006; Hammond et al., 2007) show that the dermomyotome occupies a similar domain on the surface of each somite and expresses the muscle progenitor cell marker *pax7*. Previous work in mouse and chick has shown that satellite cells originate from the embryonic dermomyotome (Gros et al., 2005; Kassar-Duchossoy et al., 2005; Relaix et al., 2005; Schienda et al., 2006). Consistent with this, using lineage tracing and time-lapse imaging, we showed that *col1a2*^*+*^ dermomyotome cells contribute to not only new muscle fibers but also small *pax7*^*+*^ fiber-associated cells, likely corresponding to larval muscle progenitor cells (Gurevich et al., 2016; Nguyen et al., 2017; Pipalia et al., 2016; Roy et al., 2017; Seger et al., 2011).

We also observed some differences between zebrafish and higher vertebrates. First, in mouse and chick, the dermomyotome has been shown to give rise to both muscles and the dermis (Christ and Scaal, 2008; Stellabotte and Devoto, 2007). Interestingly, we did not observe any contribution of *col1a2*^*+*^ dermomyotome cells to the dermis in cell tracing experiments. One possible explanation is that *col1a2*^*+*^ dermomyotome cells are already committed to the myogenic lineage by the time our *col1a2* reporters are expressed at high level in the dermomyotome at 2 dpf. Indeed, dye labeling of early dermomyotome precursors in the anterior somitic compartment prior to the somite rotation marks both muscles and some dermal-like cells (Hollway et al., 2007). Second, we found that the zebrafish dermomyotome persists in adults, similar to previous observations of *pax7* and *pax3* expression on the lateral surface of adult myotomes (Hollway et al., 2007; Nguyen et al., 2017). This persistence is in contrast with the amniotic dermomyotome, which is shown to be a transient embryonic structure (Christ and Scaal, 2008; Stellabotte and Devoto, 2007). The presence of the dermomyotome in adult zebrafish is likely required for the continuous growth of the fish.

### Unique morphology of dermomyotome cells

Using *col1a2* transgenic lines, we identified several new features of dermomyotome cells. First, dermomyotome cells display a ramified morphology. In a quiescent state, these cells are not polarized, with lamellipodia-like structures all around the cell. This morphology is in contrast with mouse satellite cells, which usually display a bipolar morphology with short processes extending along the myofiber (Webster et al., 2016). Second, although dermomyotome cell bodies are mostly stationary in uninjured conditions, their cellular processes are quite dynamic, constantly extending and retracting. Each cell and its processes occupy a largely non-overlapping area, reminiscent of the tiling of neuronal dendrites (Grueber and Sagasti, 2010). During cell division, the mother cell retracts all its processes, but immediately after the division, the two daughter cells would extend new processes to reclaim the similar surface area. These dynamic behaviors might ensure the complete coverage of the somitic surface and detect potential muscle injuries. Lastly, we often observed long filopodia-like structures extending from dermomyotome cells. Intriguingly, our single cell lineage analysis revealed that the distance between the dermomyotome cell body and the injury site is not a reliable predictor of whether the cell would generate a regenerative response. It thus raises the possibility that the long cellular projections might allow the dermomyotome cell to detect muscle injury at a significant distance away from the cell body. Consistent with our findings, dermomyotome cells in chick display similar filopodia-like protrusions, facilitating the interaction with the overlying ectoderm during somite development (Sagar et al., 2015).

### ECM genes as novel markers of dermomyotome cells

The dermomyotome is traditionally defined by muscle progenitor cell markers *pax7* and *pax3*. We demonstrated that the zebrafish dermomyotome expresses a number of ECM genes, including *col1a2*, *col5a1*, and *cilp*. Consistent with our finding, an ultrastructural expression analysis in zebrafish identifies some *col1a2*^*+*^ mesenchymal cells on the surface of muscles, likely corresponding to dermomyotome cells (Le Guellec et al., 2004). Similarly, *col1a1* and *cilp* have been shown to express in a dermomyotome-like domain in trout embryos (Ralliere et al., 2015; Rescan et al., 2005). Moreover, we found that the expression of these ECM genes is dynamically induced during muscle injury repair. Our observations in embryonic muscle progenitor cells in zebrafish show strong similarities to mouse muscle progenitor cells. For example, fetal muscle stem cells in mice show high-level expression of several ECM molecules, such as tenascin-C (*TnC*), fibronectin (*Fn1*), and collagen VI (*Col6*) (Tierney et al., 2016). Similarly, newly identified *Twist2*^*+*^ mouse muscle progenitor cells show enrichment in many ECM genes, including *Col1a1*, *Col1a2*, and *Cilp* (Liu et al., 2017). Interestingly, although quiescent adult satellite cells do not express high level of ECM genes (Liu et al., 2017), activated satellite cells show upregulation of many collagen genes, such as *Col1a1* and *Col1a2* (Pallafacchina et al., 2010). Thus, dynamic regulation of ECM gene expression might be a common feature of muscle progenitor cells.

Does the ECM play an active role in regulating the function of dermomyotome cells? We envision two non-mutually exclusive scenarios. First, the ECM might be an integral part of the niche in maintaining the self-renewing property of muscle progenitors. Recent work has implicated many ECM components, including fibronectin, tenascin-C, laminins, and collagens, as critical niche factors that modulate satellite cell function (Bentzinger et al., 2013b; Fry et al., 2017; Rayagiri et al., 2018; Tierney et al., 2016; Urciuolo et al., 2013). In particular, it has been shown that Collagen V produced by adult muscle satellite cells is an essential component of the quiescent niche, as deletion of the *Col5a1* gene results in depletion of the stem cell pool (Baghdadi et al., 2018). Second, the induction of ECM gene expression upon muscle injury might provide the appropriate scaffold for the recruitment and/or migration of activated muscle progenitor cells. Consistent with this idea, intravital imaging of muscle regeneration in mice reveals that ECM remnants from injured muscle fibers provides cues to regulate muscle progenitor cell behavior (Webster et al., 2016). It remains to be investigated whether loss of any ECM component would compromise the regenerative capacity of dermomyotome cells in zebrafish.

### Single cell dynamics of dermomyotome cells during injury repair

The existing *pax7* reporters in zebrafish label not only the dermomyotome and the overlying xanthophores, but also fiber-associated deep myotomal cells (Pipalia et al., 2016; Seger et al., 2011), which makes imaging individual dermomyotome cells challenging. To circumvent this issue, we developed a new *col1a2:Gal4* transgenic line to label and image dermomyotome cells. By taking advantage of the variegated nature of the *col1a2:Gal4; UAS:Kaede* line and the photoconvertible Kaede, we performed single cell clonal analysis of dermomyotome cells. A “quiescent” dermomyotome cell (type I/II responses) maintains its ramified morphology even after occasional cell divisions. By contrast, an “activated” dermomyotome cell (type III/IV responses) undergoes several stereotypic steps to generate new muscle fibers (Fig. 7G). First, it changes from its resting ramified morphology to an elongated and polarized morphology, usually within the first 3-5 hours after injury. The cell extends long cellular projections along the longitudinal axis of neighboring muscle fibers. Next, the “activated” cell undergoes several rounds of cell divisions generating a clone of small polarized daughter cells by 12-24 hpi. Finally, by 72 hpi, new muscle fibers emerge at the injury site, likely formed through fusion with uninjured muscle fibers (discussed below). The dynamic behaviors of an “activated” dermomyotome cell, such as the initial polarization phase, are reminiscent of *myf5*^*+*^ fiber-associated muscle progenitor cells during muscle regeneration (Gurevich et al., 2016). Interestingly, unlike *myf5*^*+*^ muscle progenitor cells, dermomyotome cells do not cross the myotendinous junction to repair muscle injury in neighboring somites. Quantification of our single cell clonal analysis further reveals that although the distance to the injury does not predict the response of the dermomyotome cell (discussed above), dermomyotome cells located along myosepta are more likely to respond to the injury than centrally located ones. This result suggests that ECM-enriched myotendinous junction might provide the scaffold to facilitate the migration of activated dermomyotome cells.

### Dermomyotome cells generate new fibers by fusion

Two independent lines of evidence indicate that dermomyotome cells generate new muscle fibers primarily through cell fusion. First, time-lapse imaging of the regeneration process in *col1a2*^*Kaede*^ embryos showed that the emergence of *Kaede*^*+*^ muscle fiber always appeared to be completed between two time points (8-minute interval), accompanied by the simultaneous disappearance of the dermomyotome cell. The absence of intermediate steps, such as searching for attachment sites along the myotendinous junction, suggests that new fibers are formed by cell fusion rather than *de novo* fiber formation. Second, our Cre-mediated lineage tracing experiments showed that most new muscle fibers express both EGFP and mCherry, indicating cell fusion events between a switched dermomyotome cell (*mCherry*^*+*^, *ubi:loxP-mCherry*) and an un-switched muscle fiber (*EGFP*^*+*^, *ubi:loxP-EGFP-loxP-mCherry*). Intriguingly, during muscle injury repair, significantly more *de novo* fiber formation (14%) was observed compared to uninjured conditions (3%). This discrepancy might be explained by differential activation of sub-populations of dermomyotome cells. Recent work shows that two paralogues of *pax7*, *pax7a* and *pax7b*, mark similar but not identical muscle progenitor cell populations. *pax7b*^+^ cells contribute to muscle growth and repair by cell fusions, whereas *pax7a*^*+*^*pax7b*^*-*^ cells predominantly generate nascent fibers (Pipalia et al., 2016). Our results suggest that normal muscle growth is primarily contributed by *pax7b*^*+*^ dermomyotome cells, while muscle injury activates both *pax7b*^+^ and *pax7a*^*+*^*pax7b*^*-*^ cells. Consistent with this idea, *pax7a*^*+*^ cells have been shown to contribute to only large muscle injuries but not small injuries (Knappe et al., 2015).

In summary, our work provides a dynamic view of dermomyotome cells during muscle growth and repair. It also raises additional questions for future investigations. For example, what is the injury signal that activates dermomyotome cells? Tissue injury is often associated with elevated level of reactive oxygen species (ROS) and the recruitment of patrolling immune cells such as macrophages and neutrophils (Love et al., 2013; Niethammer et al., 2009). It is therefore plausible that ROS and/or cytokines secreted by immune cells might be the trigger to activate dermomyotome cells. Indeed, mice and human data have implicated macrophages and other immune cells as the critical regulators of satellite cell functions (Bentzinger et al., 2013a; Saclier et al., 2013).

## MATERIALS AND METHODS

### Zebrafish strains

Zebrafish strains were maintained and raised according to the standard protocols (Westerfield, 2000). All procedures were approved by the University of Calgary Animal Care Committee. Embryos were grown at 28.5 °C and staged as previously described (Kimmel et al., 1995). Fish older than 24 hpf were treated with 1-phenyl 2-thiourea (PTU) to prevent pigmentation. TL and TL/AB wild-type strains were used in this study along with the following transgenic lines: *α-actin:GFP* (Higashijima et al., 1997), *col1a2:Gal4*, *UAS:Kaede* (Scott et al., 2007), *UAS:NTR-mCherry* (Davison et al., 2007), *UAS:Cre-ERT2*, *Ubi:loxP-EGFP-loxP-mCherry* (*Ubi:Switch*) (Mosimann et al., 2011). The mosaic *col1a2:Gal4; UAS:Kaede* line was maintained by selectively growing embryos with more mosaic Kaede expression.

### Generation of transgenic lines

*UAS:Cre-ERT2* was generated by standard Tol2-mediated transgenesis. To generate *col1a2:Gal4* transgenic line, BAC clone zC122K13 from the CHORI-211 library that contains *col1a2* genomic region with 78 kb upstream and 58 kb downstream regulatory sequences was selected for bacteria mediated homologous recombination following the standard protocol (Bussmann and Schulte-Merker, 2011). Briefly, the pRedET plasmid was first transformed into BAC-containing bacteria. Second, an iTol2_amp cassette containing two Tol2 arms in opposite directions flanking an ampicillin resistance gene was recombined into the vector backbone of zC122K13. Lastly, a cassette containing the Gal4-VP16 with a kanamycin resistant gene was recombined into zC122K13-iTol2_amp to replace the first coding exon of the *col1a2* gene. After each round of recombination, successful recombinants were confirmed by PCR analysis. The final *col1a2:Gal4* BAC was then co-injected with *tol2* transposase mRNA into *UAS:Kaede* embryos at one-cell stage. Positive transgenic lines were identified by screening Kaede expression in F1 embryos from injected founders.

### Muscle injury

Two methods were employed to generate muscle injury at specific locations in larval zebrafish. In needle injury, we used a sharp injection needle to stab muscles near the end of yolk extension (somites 17-19) so the injury site can be recognized easily during the lineage tracing. Alternatively, to introduce muscle injury at a more precise location, laser ablation was performed with the 750 nm laser and the 25x objective on the Leica TCS SP8 multi-photon microscope. A region of interest (ROI) at a desired location was selected, zoomed in to the maximum (48x), and scanned with 100% 750 nm laser once. The laser-induced injury can be readily visualized in the bright field after the scanning.

### Kaede photoconversion

*col1a2*^*Kaede*^ embryos at appropriate stages were anesthetized with tricaine and mounted in 0.8% low melting point agarose in a glass bottom dish (MatTek). Photoconversion experiments were performed using the 405 nm laser and the 20x objective on the Olympus FV1200 confocal microscope. For photoconversions of large areas (∼ 5 to 6 somite region), 50% laser power was used to scan the desired ROI for 2 frames at a dwell time of 200 μs per pixel. For single cell photoconversions, 2% laser power was used to scan a small ROI (10 x 10 pixels) with the Tornado mode at a dwell time of 2 μs per pixel for a total of 1-2 seconds. After photoconversion, embryos were released from the agarose, transferred to fish water to recover in the dark, and analyzed at desired stages.

### Cre-mediated lineage tracing

To obtain mosaic labeling, *col1a2:Gal4; UAS:Cre-ERT2; ubi:Switch* embryos were pulsed with 10 μM 4-hydroxytamoxifen (4-OHT) for 2-3 hours at desired stages. After treatment, 4-OHT was washed off with fish water for three times, and embryos were recovered in fish water for analysis at appropriate stages.

### Single cell lineage tracing

Mosaic *col1a2*^*Kaede*^ embryos at appropriate stages were selected for single cell tracing experiments. Individual isolated cells were photoconverted to *Kaede*^*red*^. To ensure reliable cell tracing over time, a maximum of one cell per somite and four cells per embryo were photoconverted and traced. Immediately after photoconversion, muscle injury was introduced by laser ablation near one *Kaede*^*red*^ cell. Images were taken before and after the photoconversion and then every 24 hours till 72 hpi to trace individual *Kaede*^*red*^ cells. For quantification in Fig. 7D, distance was measured from the center of the photoconverted cell to the center of the muscle injury.

### In situ hybridization and immunohistochemistry

Whole-mount in situ hybridization and antibody staining were performed according to the standard protocols (Thisse et al., 2004). The following antisense probes were used in this study: *cilp*, *col1a1a*, *col1a2*, *col5a1*, *kaede*, *myoD*, *myogenin*, *pax7* (Seo et al., 1998), *postnb* and *sparc*. For antibody staining, the following primary antibodies were used: rabbit polyclonal antibody to Kaede (1:1000, MBL) and mouse monoclonal antibody to Pax7 (1:10, Developmental Studies Hybridoma Bank (DSHB)). For fluorescent detection of antibody labeling, appropriate Alexa Fluor-conjugated secondary antibodies (1:500, Molecular Probes) were used.

### Cell ablation experiments

To ablate *col1a2*^*+*^ cells, *col1a2*^*NTR-mCherry*^ transgenic fish were outcrossed with wild-type fish to obtain *mCherry*^*+*^ embryos (experimental group) and *mCherry*^*-*^ embryos (control group). Embryos at 48 hpf were treated with metronidazole (MTZ) at a final concentration of 5 mM in fish water for 24 hours. Embryos were then washed with E3 fish water 2-3 times and grown to desired stages for analysis.

### Time-lapse imaging and processing

Embryos were anesthetized with tricaine and embedded in 0.8% low melting point agarose on a glass bottom dish (MatTek). Fish were imaged with Olympus FV1200 confocal microscope using the 20x objective. For time-lapse imaging of muscle injury repair, embryos were first injured at 3 dpf either by needle stabbing or laser ablation. Injured embryos were then imaged laterally starting at either 0 or 29 hpi at 8-min intervals for 19-20 hours. All the confocal images were analyzed and quantified using the Fiji software (Schindelin et al., 2012). Brightness and contrast were adjusted for better visualization. To generate color-coded depth projections, confocal z-stacks were processed with Fiji using the ‘temporal color code’ function.

### Quantification of dermomyotome cells

*col1a2*^*NTR-mCherry*^ fish were stained with anti-Pax7 antibody at 2 dpf. Cells in each somite were counted based on *col1a2* expression and Pax7 staining: all Pax7^+^ cells, Pax7^+^Col1a2^+^ cells and Pax7^+^Col1a2^-^ cells. Note that xanthophores, which have strong Pax7 staining, were not counted. Overall, average number of cells in five somites per embryo was used for the quantification. Pax7^+^Col1a2^+^ cells were further categorized according to their position in a somite: near the vertical myosepta (VM), near the horizontal myoseptum (HM), or in between somatic boundaries (central).

### Data analysis

All the graphs were generated in the GraphPad Prism software. For quantifications, standard error of the mean was calculated. To analyze significance between two samples, P values were determined by performing the Mann-Whitney *U* test.

## ACKNOWLEDGEMENTS

We thank the zebrafish community for providing probes and reagents; Holger Knaut for BAC clones; Jason Berman for *UAS:NTR-mCherry* fish; Sarah Childs and members of the Huang laboratory for discussion; and Paul Mains and James McGhee for critical comments on the manuscript.

## SUPPLEMENTARY INFORMATION

**Figure S1.**
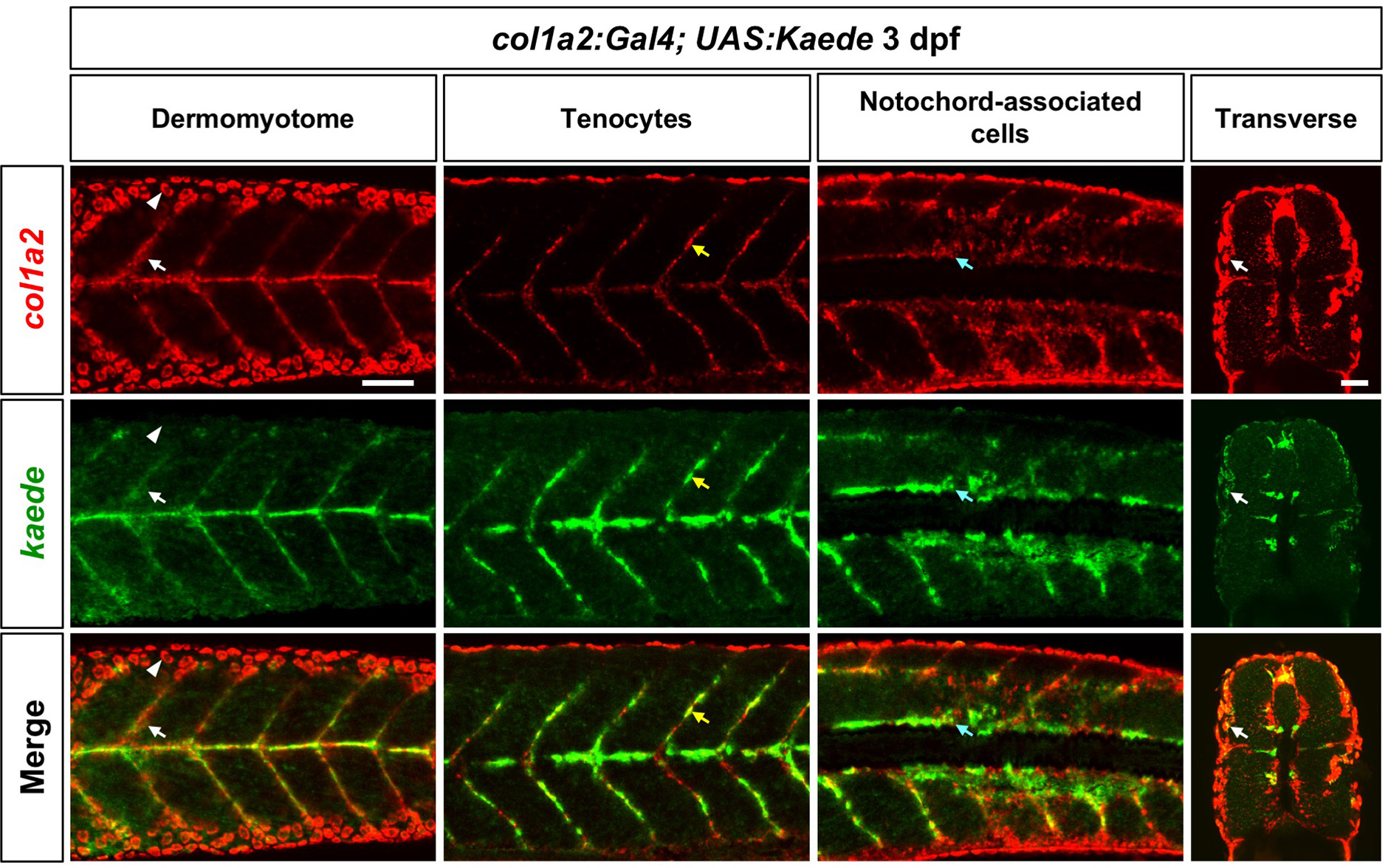
Validation of the ***col1a2*^*Kaede*^ line**. Double fluorescent in situ hybridization using *kaede* and *col1a2* probes were performed in *col1a2*^*Kaede*^ embryos at 3 dpf. Co-expression of *kaede* (green) and the endogenous *col1a2* (red) can be observed in dermomyotome cells (white arrows), tenocytes along the vertical myoseptum (yellow arrows), and deep interstitial cells around the notochord (cyan arrows). Note that *col1a2*^*Kaede*^ was not expressed in the skin cells (arrowheads) as *col1a2*. Scale bar: 50 µm.

**Figure S2.**
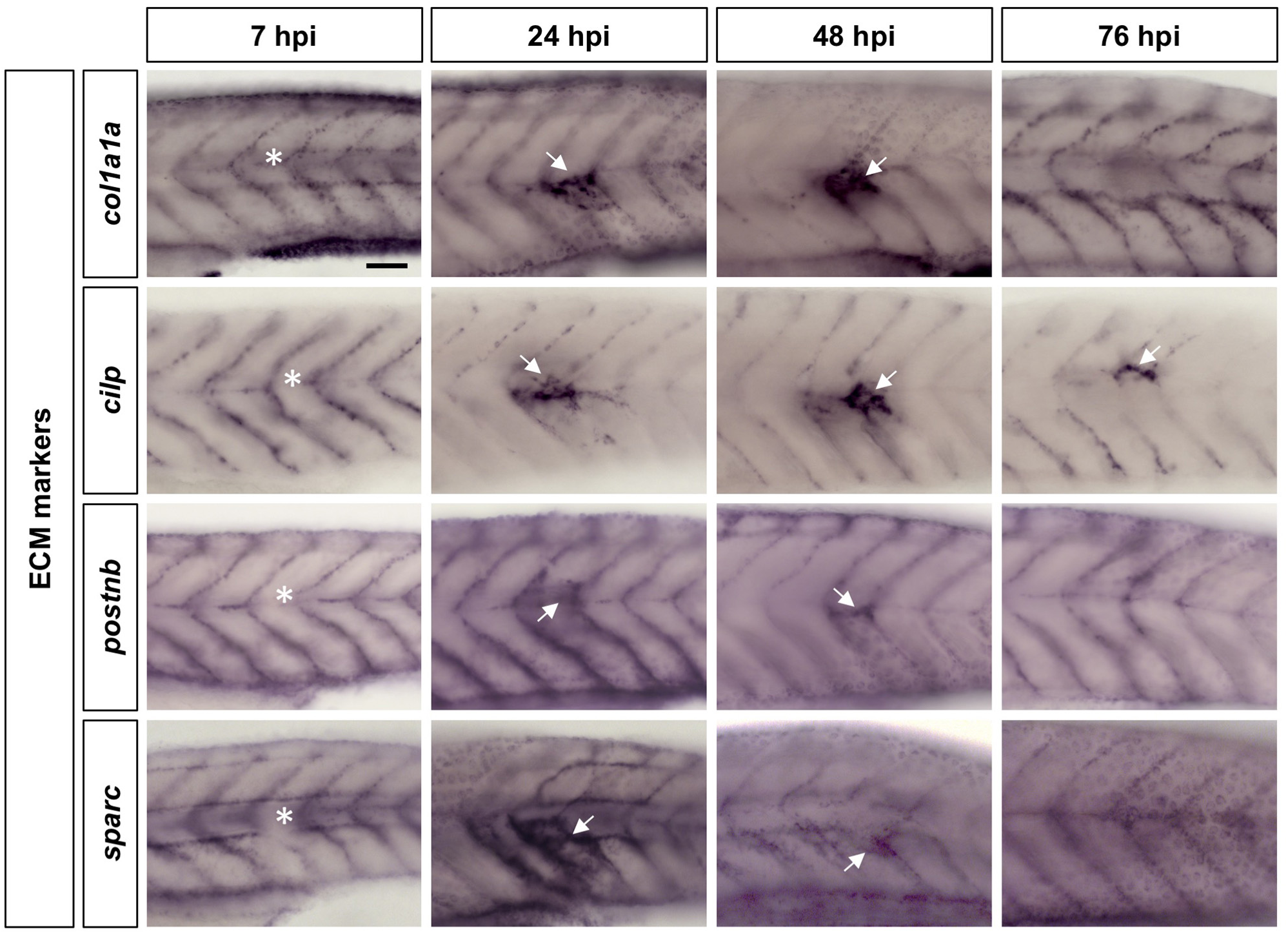
ECM molecules are upregulated during muscle injury repair. Wild-type embryos were needle stabbed to injure a somite near the end of yolk extension (asterisks) at 3dpf, and fixed at 7, 24, 48, and 76 hpi. Embryos were then stained with ECM markers (*col1a1a*, *cilp*, *postnb*, and *sparc*). All markers showed upregulation at the site of injury starting from 24 hpi (arrows). Scale bars: 50 µm.

**Figure S3.**
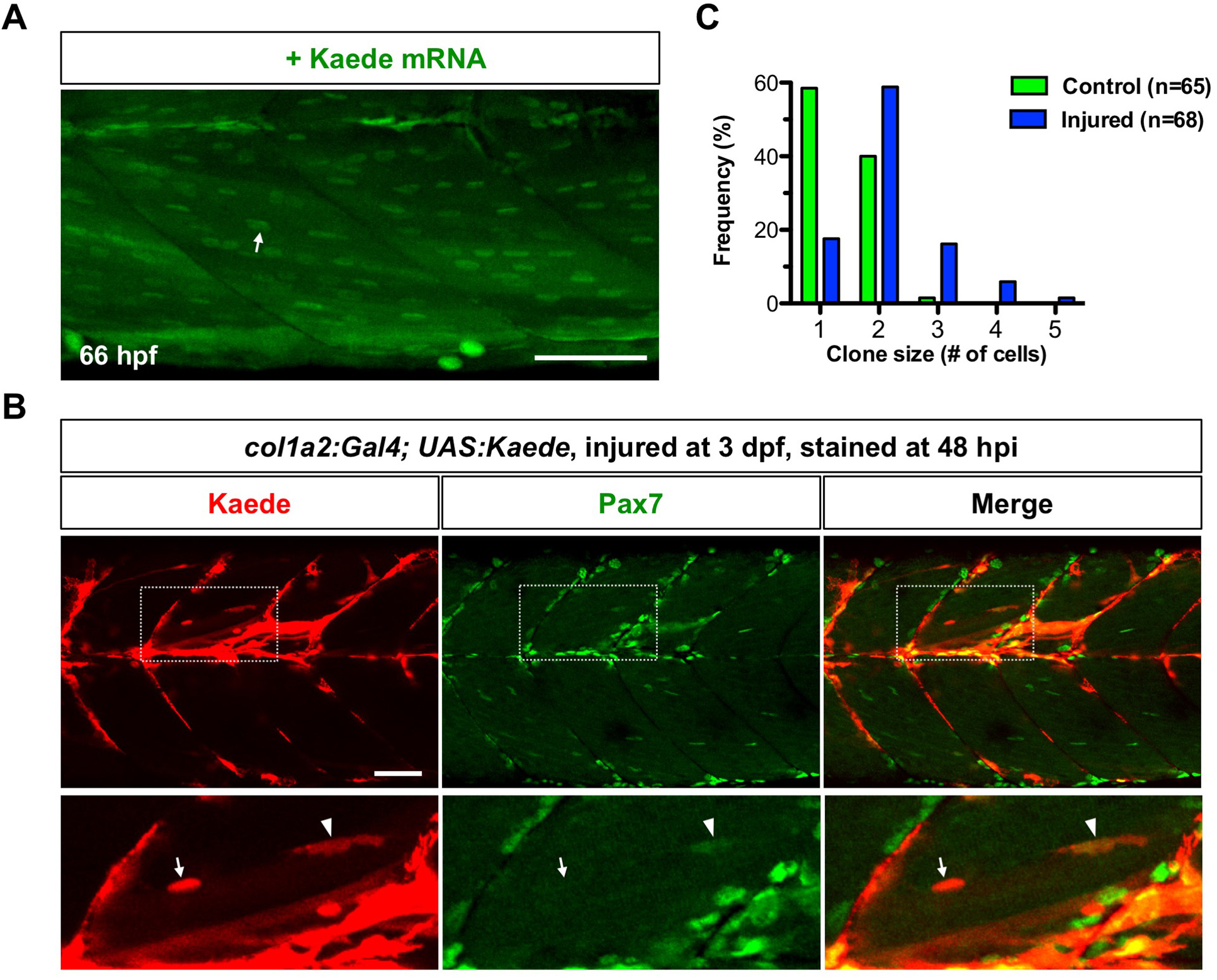
Kaede protein is preferentially localized in nuclei of muscle fibers. (A) Wild-type embryos were injected with Kaede mRNA, and imaged at 66 hpf. Kaede protein (green) is preferentially localized in the nuclei of muscle fibers (arrow). (B) *col1a2*^*Kaede*^ embryos were injured at 3 dpf, and stained at 48 hpi with the anti-Pax7 antibody (green). *Kaede*^*+*^ dermomyotome derived cells (red) contributed to muscle injury repair (boxed regions). The expanded views show that newly formed muscle fiber displayed strong Kaede expression in the nucleus (arrows), which was not labelled by Pax7. By contrast, a small elongated *Kaede*^*+*^ cell between muscle fibers was Pax7 positive (arrowheads). (C) Quantification of clone size in single cell clonal analysis described in Fig 5. Dermomyotome cells under the injury condition (blue, n=68) tend to generate larger clones compared to cells in the wild-type condition (green, n=65). Scale bars: 50 µm.

**Movie 1. Expression pattern of the *col1a2* transgenic line.** A confocal z-stack of *col1a2:Gal4; UAS:NTR-mCherry; α-actin:GFP* embryos at 3 dpf shows mCherry expression (green) in dermomyotome cells, some muscle fibers, tenocytes and notochord-associated cells (arrows). Muscle fibers are labeled with *α-actin:GFP* (magenta). Scale bar: 50 *µ*m.

**Movie 2. Dynamics *col1a2*^*+*^ dermomyotome cells in quiescent state.** *col1a2:Gal4; UAS:NTR-mCherry; UAS:Kaede* embryos were imaged at 2 dpf for 7.9 hours. *col1a2*^*+*^ dermomyotome cells cover the entire surface of a somite. When a dermomyotome cell divides, daughter cells reclaim the same surface area soon after division. Five different cell divisions are indicated by arrows. Scale bars: 50 *µ*m.

**Movie 3. Dynamics *col1a2*^*+*^ dermomyotome cells in injured condition.** *col1a2*^*Kaede*^ embryos were injured, photoconverted at 59 hpf, and then imaged over 23 hours (0-23 hpi). Cells “a” and “b” were within the injured area while the cell “c” was in the uninjured area. Cell “a” (white arrows) maintained the ramified morphology, and divided once at 7 hpi generating two daughter cells with similar morphologies. By contrast, cell “b” (yellow arrows) extended to form an elongated morphology (arrowheads), and divided once at 16 hpi generating two polarized daughter cells. Cell “c” (cyan arrows) did not divide and remained its ramified morphology. Scale bars: 50 *µ*m.

**Movie 4. Generation of new muscle fibers by cell fusion.** *col1a2*^*Kaede*^ fish was injured at 3 dpf and imaged from 29 hpi onwards. A *Kaede*^*+*^ dermomyotome cell (arrows) near the injury site elongated at 34 hpi (white arrowheads), formed protrusions at 39 hpi, and fused with a neighboring muscle fiber at 40 hpi. The newly formed muscle fiber can be identified by the weak Kaede expression throughout the muscle fiber and the strong Kaede expression in the nucleus (yellow arrowheads). Scale bars: 50 *µ*m.

